# Premovement suppression of corticospinal excitability may be a necessary part of movement preparation

**DOI:** 10.1101/470153

**Authors:** J. Ibáñez, R. Hannah, L. Rocchi, J.C. Rothwell

## Abstract

In a warned reaction time (RT) task, corticospinal excitability (CSE) decreases in task-related muscles at the time of the imperative signal (preparatory inhibition). Because RT tasks emphasise speed of response, it is impossible to distinguish whether preparatory inhibition reflects a mechanism preventing premature reactions, or whether it is an inherent part of movement preparation. We used transcranial magnetic stimulation (TMS) to study CSE changes preceding RT movements and movements that were either self-paced (SP) or performed at predictable times to coincide with an external event (PT). Results show that CSE changes over a similar temporal profile in all cases, suggesting that preparatory inhibition is a necessary state in planned movements allowing the transition between rest and movement. Additionally, TMS given shortly before the times to move speeded the onset of movements in both RT and SP contexts, suggesting that their initiation depends on a form of trigger that can be conditioned by external signals. On the contrary, PT movements do not show this effect, suggesting the use of a mechanistically different triggering strategy. This relative immunity of PT tasks to be biased by external events may reflect a mechanism that ensures priority of internal predictive signals to trigger movement onset.

## Introduction

A variety of different forms of inhibition related to volitional movement have been observed in the motor system, such as reactive inhibition, selective inhibition and proactive inhibition (Duque et al. 2017). However, one of the least understood is preparatory inhibition, which describes markers of inhibition that are observed prior to the onset of a voluntary movement (Hasbroucq et al. 1997; Duque and Ivry 2009). Preparatory inhibition has been investigated intensively using TMS to monitor excitability of the corticospinal output to muscles that are both involved or not involved in the task (Duque and Ivry 2009; Duque et al. 2010; Bestmann and Duque 2015; Greenhouse et al. 2015). For example, in a warned reaction time task, corticospinal excitability (CSE) is depressed in both the selected and non-selected effectors just prior to the onset of the imperative signal (Hasbroucq et al. 1997; Touge et al. 1998). Three explanations have been proposed for this inhibition. Competition resolution proposes that inhibition is necessary to suppress competing movements, at least in situations in which the selected movement is not precisely known in advance (Burle et al. 2004); a second possibility is that inhibition is necessary to prevent premature release of a prepared, “subthreshold” movement (Duque and Ivry 2009; Duque et al. 2010); a third possibility, sometimes known as the spotlight hypothesis, is that preparatory inhibition reduces background motor activity. This speeds movement onset because excitatory inputs that select the chosen response stand out better against a quiescent background (Greenhouse et al. 2015; Lebon et al. 2018).

In a recent study, Hannah et al. argued against the premature release hypothesis. In single trials they found that the greater the corticospinal suppression to the selected effector, the faster the reaction time, which was more compatible with the idea that inhibition was facilitating movement rather than suppressing it (Hannah et al. 2018). It was more consistent with the “spotlight/noise reduction” hypothesis than prevention of release. However, they made another observation that seemed inconsistent with noise reduction. Suppression was only observed with MEPs evoked by an antero-posterior monophasic current pulse, but was much smaller or absent with a postero-anterior pulse. The two directions of current pulse activate two different sets of synaptic inputs to corticospinal output neurones (Hamada et al. 2014; Hannah and Rothwell 2017). A simple version of noise reduction should predict reduced activity in both these pathways rather than the selective effect they reported. Because of this they proposed that the results were consistent with recent work of Churchland et al. in primates that proposes that preparation for a forthcoming movement can occur without incurring change in corticospinal output because it is prepared in a neural activity space that is orthogonal to that employed in the ongoing task (Churchland et al. 2010; Kaufman et al. 2014). Since both activity spaces converge on the same corticospinal output neurones, this implies that although some inputs to corticospinal neurones are more active during preparation, others are less active or inhibited. Hannah et al suggested, but without explaining how, that their PA pulse preferentially tested those inputs that were suppressed.

All previous TMS experiments studying preparatory inhibition have involved some version of a reaction time (RT) task. The aim of the present experiments was to expand the study of preparatory inhibition to other types of movement. Specifically, we compared preparatory inhibition in RT movements with that obtained in two other movement types: (1) a predictive task (PT) in which movement initiation is timed to coincide with the last event in a countdown sequence, and (2) in self-paced (SP) movements that are devoid of any external trigger. The same movement was required in all three paradigms. Importantly, it had no choice element so that participants knew in advance exactly what was required on each trial. This removes the necessity for conflict resolution (see also Quoilin et al. 2019) and focuses the question on whether preparatory inhibition is preventing (premature release) or aiding movement (“spotlight hypothesis” or “orthogonal activity” as suggested by Hannah et al (2018)).

Finally, because we used single-pulse TMS to probe CSE, we were also able to address a second feature typical of movements in RT tasks: intersensory facilitation (IF), i.e., the significant reduction of reaction times and the speeded release of the prepared movements when a secondary stimulus (a TMS pulse in our case) is delivered at about the time of the imperative stimulus (Nickerson 1973). In movements made in a RT context, IF is usually explained in terms of shortening the time taken to identify the imperative stimulus (Pascual-Leone, Valls-Sole, et al. 1992). SP and predictive movements may not require an external trigger since they can start immediately once preparation is complete. If this is the case, they should not display IF.

## Materials and Methods

### Participants

In total, 33 right-handed healthy subjects participated in this study (28 ± 1 years old; age range 20-45 years; 15 females). Fifteen participants (7 females) took part in experiment 1 and 18 (8 females) took part in experiment 2. All of them reported no contraindications to TMS (Rossi et al. 2011) and had normal or corrected to normal visual acuity. The study was approved by the University College London Ethics Committee and warranted to be in accordance with the Declaration of Helsinki. All participants signed a written informed consent prior to the experimental session.

### Recordings

Participants sat in a comfortable chair with both forearms resting on a pillow placed on their lap and index fingers resting on a keypad through which button press times were recorded. A screen was placed ~1 m in front of the participants. They also wore ear defenders to reduce the influence of loud sounds generated by the TMS discharges.

EMG signals were obtained from the right first dorsal interosseous (FDI) muscle for experiment 1 and of both hands in experiment 2. EMG activity from the right abductor digiti minimi (ADM) muscle was also recorded. MEPs amplitudes recorded from the right FDI were the primary dependent measure in this study. The ADM was used as a control muscle for the assessment of MEP changes: it was activated by the TMS but, unlike the right FDI, it was not directly involved in the response, so it was only supposed to show a monotonic reduction of excitability up until the time at which muscles showed voluntary activation (Duque et al. 2010). Recording electrodes were placed on the muscle bellies, with reference electrodes on the closest metacarpophalangeal joint. The ground electrode was placed on the right wrist. EMG signals were amplified, band-pass filtered between 20 Hz and 2000 Hz (Digitimer D360, 2015 Digitimer Ltd, United Kingdom) and acquired at 5000 Hz sampling rate with a data acquisition board (CED-1401, Cambridge Electronic Design Ltd 2016) connected to a PC and controlled with either Signal or Spike^2^ software (also by CED).

Once EMG sensors were set, the participants’ TMS hotspot was located. This was done by finding the point over M1 giving the largest MEPs in the contralateral FDI for a given stimulus intensity. The TMS coil was held at a 45° angle to the sagittal plane with the handle pointing backwards. Once the hotspot was found, the resting motor threshold (RMT) and 1 mV intensity were determined. The RMT was estimated by adjusting the TMS output until 5 out of 10 MEPs larger than 50 μV could be obtained. The 1 mV intensity was defined by adjusting the TMS output until 5 out of 10 MEPs larger than 1 mV could be obtained. The estimated 1 mV TMS intensity was then used to assess CS excitability changes in the three paradigms tested in this study.

The experimental paradigms were implemented using custom-made MATLAB routines (MathWorks, MA, USA). Synchronization of TMS pulses with EMG and movement events was realised using Cogent 2000’s utilities (Cogent 2000 team at the FIL and the ICN and Cogent Graphics developed by John Romaya at the Wellcome Department of Imaging Neuroscience) to control the parallel port of the PC running the experimental paradigms. Data analysis was carried out using custom-made MATLAB functions and SPSS software (IBM, NY, USA).

### Experiment 1 – TMS recordings preceding movements in RT and PT tasks

For this experiment, participants performed two types of movement paradigms: 1) a RT task in which movements were initiated following an imperative stimulus (Fig. 1-A); and 2) a PT task in which movements were timed with an external countdown signal (Fig. 1-B). In both cases, each trial of the motor task consisted in pressing a button of a keypad with the right index finger at the times indicated by a visual cue. Unilateral movements were chosen to make experiments as similar as possible to most simple reaction time tasks used in previous TMS studies on movement preparation (Duque et al. 2010; Greenhouse et al. 2015; Hannah et al. 2018).

**Figure 1.**
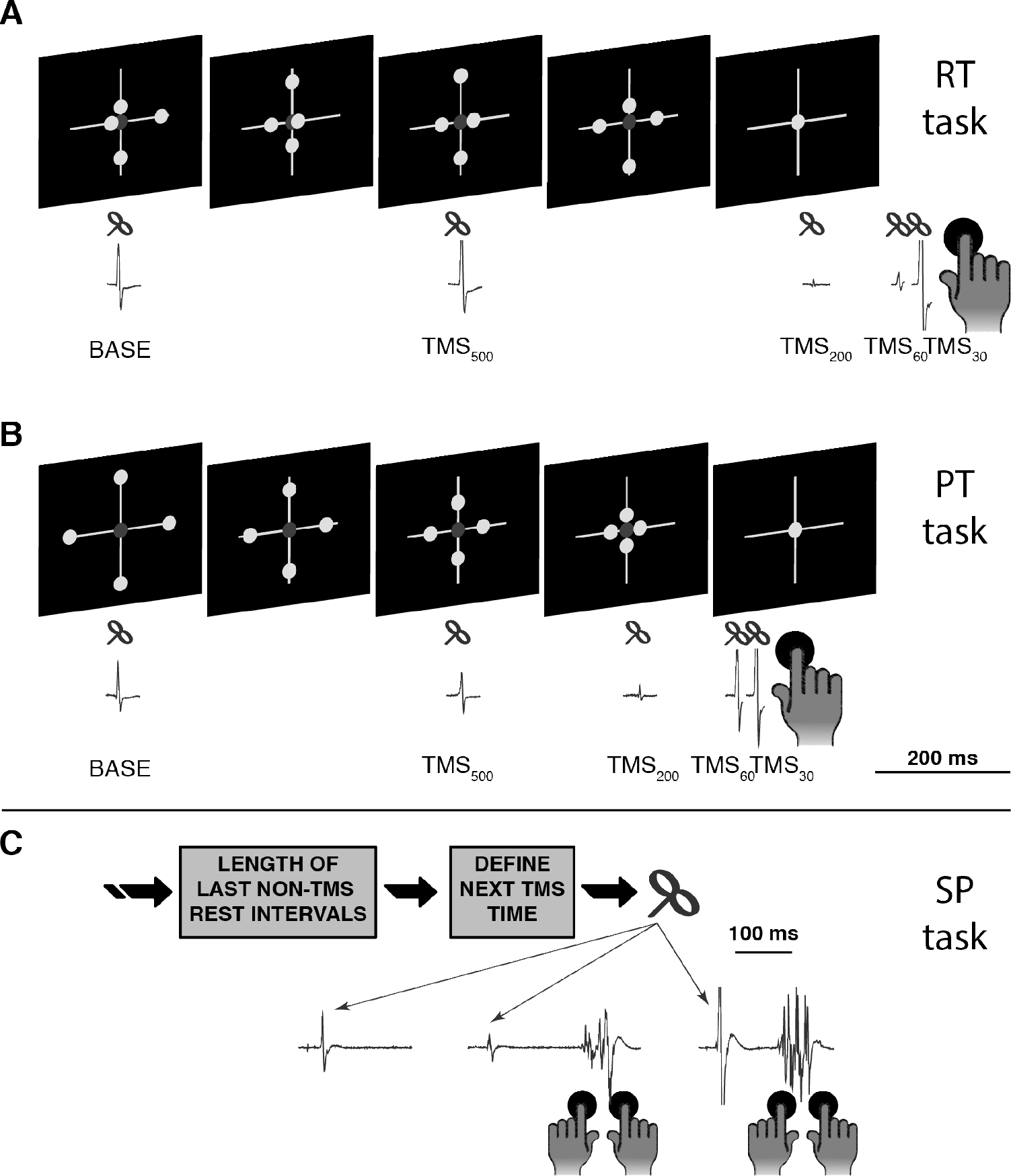
Movement tasks and TMS recordings. (A) In each trial of the reaction time (RT) task -experiment 1- four circles move randomly along the axes of a cross for 1 s. After this delay period, all circles suddenly collapse at the intersection point of the cross and this is the “GO” signal making participants react as fast as they can, performing a button press with their right index finger. (B) In the predictive time (PT) task -experiment 1- participants have to perform the button press at the end of a 1-s countdown period, which is informed by showing four white circles moving along the axes of a cross reaching simultaneously the intersection point from the extremes. Both in PT and RT paradigms, single-pulse TMS was either not delivered (non-TMS trials) or delivered when the four circles appeared onscreen at the beginning of the delay period (BASE), half way through the delay period (TMS_500_), and 200 ms (TMS_200_), 60 ms (TMS_60_) and 30 ms (TMS_30_) before the average EMG onset time of each participant in each task. (C) In experiment 2, participants performed self-paced (SP) movements consisting of simultaneously pressing two keypad buttons with the index fingers of their two hands. An algorithm was run in parallel to characterize the times at which movements were performed in the non-TMS trials. This information was in turn used to distribute TMS pulses in subsequent TMS trials with different time intervals between the stimuli and the movements.

**Figure 2.**
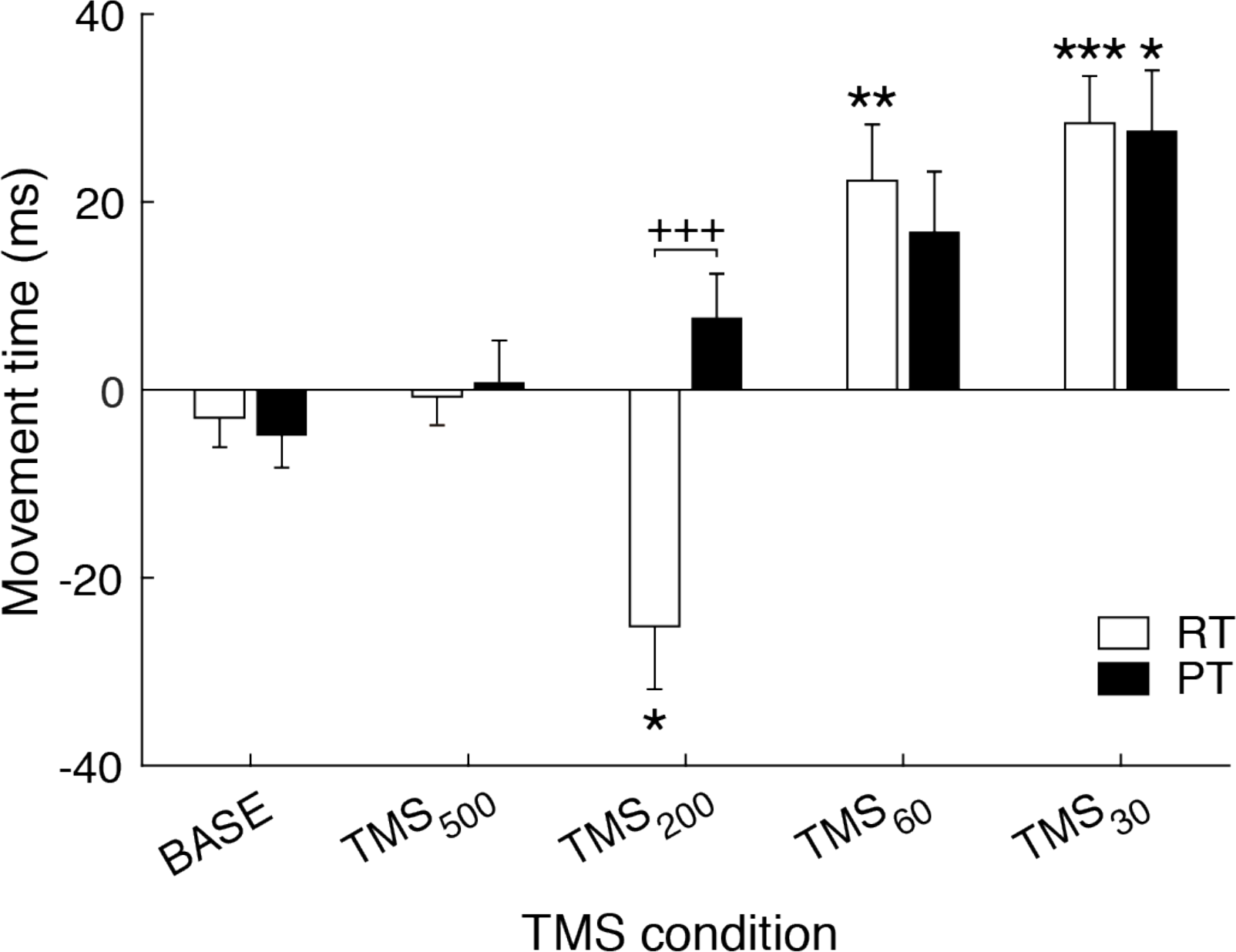
Movement times. (relative to non-TMS trials) for the different TMS conditions and for the RT (white) and PT (black) tasks. \**P* < 0.05; \*\**P* < 0.01; \*\*\**P* < 0.001, compared with BASE time point within each movement task. +++*P* < 0.001, RT vs PT.

Each trial of the RT task consisted of a resting phase of 2 s followed by a delay period of 1 s during which four circles moved randomly along the four arms of a cross. After the end of the delay period, the four circles were plotted at the intersection point of the cross, and this event was to be considered the “GO” cue. Participants were instructed to make a fast and ballistic button press with their right index finger in reaction to seeing the “GO” cue. After each button press, the trial ended by giving participants feedback about the time at which the button press event had been detected. In the set-up used in these experiments, an interval of ~90 ms separated the FDI EMG onset and the time at which the button press event was detected. The feedback was displayed for a random period of time between 1-3 s, and it consisted of the time at which the button press had been detected and a font colour code indicative of the performance. Button presses in the interval 250-300 ms resulted in giving feedback about the time of the response with green text (presses within this interval implied that the FDI activation onset had taken place with a reaction time of 160-210 ms in most cases). Yellow text was used for button presses in the intervals 200-250 ms and 300-350 ms. Finally, red text and the warning messages “too early” and “too late” were given as feedback in case button presses were performed before or after these intervals.

The PT task had the same structure of trials as the RT task but, in this case, during the delay period the four circles moved from the extremes of a cross towards its centre with a velocity inversely proportional to the remaining distance to the intersection point (initial distance 4.5 cm). Participants were instructed to time their movements with the overlapping of the four circles at the intersection point. Since PT movements were supposed to be performed at around the time at which the circles collapsed, the feedback was different from the one used in the RT task. Green text was used for button presses done between 50-100 ms relative to the time at which circles overlapped, thus encouraging participants to aim at pressing the button within this interval (which in turn implied activating the FDI muscle at around the time at which circles collapsed at the intersection point of the cross); yellow text was used for button presses in the intervals 0-50 ms and 100-150 ms; red text was used in any other case. Additionally, the messages “too early” and “too late” were displayed when button presses were performed before or 150 ms after circles collapsed.

In the initial part of each experiment, participants practised the RT and PT paradigms without TMS until they showed consistent response times (~30 trials per task). Thirty additional movements were then performed for each paradigm so that the subject- and task-specific average movement onset times based on the FDI EMG activity (EMG onset times) could be estimated. After the initial training phase, the TMS recordings were carried out, consisting in two blocks of 78 trials per paradigm. The blocks of the two paradigms were interleaved. In each block, six conditions were tested using a randomized order of TMS conditions. TMS conditions differed from each other with regards to the timing of the stimulus: 1) no TMS delivered (non-TMS condition); 2) TMS at the beginning of the delay period (BASE); 3) TMS halfway through the delay period (TMS_500_); and 4-5-6) TMS 200 ms (TMS_200_), 60 ms (TMS_60_) and 30 ms (TMS_30_) before the average EMG onset time, respectively. Feedback was omitted in TMS trials to avoid that participants tried to compensate the possible influence that TMS could have on EMG onset times (Pascual-leone et al. 1992; Terao et al. 1997; Ziemann et al. 1997).

### Experiment 2 – TMS during the resting phases between SP movements

The task involved participants sitting still and comfortably, with both index fingers resting on a keypad. They were instructed to make ballistic bilateral button presses with the left and right hand index fingers every 4-8 s, whilst avoiding pre-movement muscle activation and ensuring movements were always made in a similar way (Fig. 1-C). Bilateral movements allowed us to measure EMG onset times from the left (non-stimulated) hand to estimate the intervals between the TMS pulses and subsequent movements without being affected by the TMS-induced delays of EMG voluntary activations of the right hand FDI in cases where the stimulus was given in close proximity with the intended EMG onset time (Ziemann et al. 1997). Importantly, bilateral synchronous actions present almost identical EMG onset times when no stimulus is given (Schneider 2004). Participants were instructed to perform their movements spontaneously and to avoid any form of internal countdown to decide when to initiate the movements. It was stressed to participants that they must not let the TMS alter their decision to move. A resting period of time followed by a button press was considered a trial, and 12 blocks were performed by each participant with 65 trials making up a block. During blocks, EMG was monitored to ensure the hand was relaxed between button presses.

A custom-made MATLAB program was used to determine the timing of a TMS stimulus on a given trial based on the timings of the button presses in the previous 5 trials performed by each participant without TMS (this number of trials was empirically chosen to allow the code program to quickly adapt to changes in participants’ behaviours). TMS timing was distributed so that in 4% of the trials, stimuli were delivered early after the previous movement (3 s after the previous button press); 8% of the trials were non-TMS trials, which were then used to monitor inter-movement intervals in the absence of external stimuli along the experiment. Finally, in 82% of the trials, TMS pulse timings were defined based on the probability density function of inter-movement intervals considering the 5 most recent non-TMS trials. For that, a Gaussian fit was estimated and the next TMS firing time was selected according to the left-hand side of this probability density function. TMS firing times were thus programmed to be delivered at a time interval relative to the previous button press such that it was always below the average inter-movement interval estimated. In the cases when participants waited for over 10 s between button presses, participants were given an indication by the experimenter to reduce the inter-movement time intervals.

### Data processing and statistical analysis

In both experiments, the onsets of the EMG were used as the reference points indicating the times of movement initiations. In order to obtain these EMG onset times in each trial, EMG was first rectified and then a moving average of 5 ms was applied to obtain a smoothed envelope of the EMG signal. EMG recordings of all trials whilst participants were at rest were analysed to obtain subject-specific resting EMG levels. Thresholds set at five times these levels were used to determine EMG onset times. These levels were also used to detect and remove trials with pre-TMS or pre-movement activation of all the muscles registered. All trials were then visually inspected and manually corrected to ensure that EMG-based movement onsets were estimated properly and that no building-up of EMG activity was apparent before the TMS. Two repeated measures ANOVA (rmANOVA) tests were run separately with data from experiments 1 and 2 to assess that EMG peak-to-peak amplitudes in the right FDI during the 200 ms intervals preceding the TMS pulses did not differ across the TMS conditions tested, *i.e.*, five TMS times tested in experiment 1 and three TMS times in experiment 2 (see below and results section). Results of these tests showed that the background EMG activity in the right FDI was not significantly different across conditions (*P* > 0.2 in all cases). Finally, EMG onset times and peak-to-peak amplitudes of MEPs were estimated. MEP amplitudes were estimated from the acquired EMG signals without applying any additional filters. A logarithmic transformation of MEP amplitudes was performed before the statistical tests to ensure normality of the samples.

To select the tests to compare EMG onset times and log-transformed MEP amplitudes across paradigms, TMS conditions and muscles in experiments 1 and 2, normality was checked by assessing that z-scores of the populations’ kurtosis and skewness were below a critical value of 2 (Kim 2013). All compared variables extracted from the TMS recording blocks satisfied the condition of normality. To compare EMG onset times between the training trials and non-TMS trials in TMS blocks in experiment 1, two Wilcoxon signed rank tests (one for each movement paradigm) were used given that normality (assessed using the Shapiro-Wilk test in this case due to the reduced length of the compared samples) was not satisfied. Finally, to run multiple comparisons of the intervals between TMS times and EMG onset times in experiment 2, both t-test and Wilcoxon signed rank tests were run. Since the two tests returned equivalent outcomes, results from the Wilcoxon analysis are shown here.

To compare EMG onset times and MEP amplitudes across conditions in experiment 1, log-transformed MEP amplitudes and times of EMG (right FDI) onsets of all trials were labelled according to the type of paradigm (PT, RT) and to the time at which TMS was delivered (BASE, TMS_500_, TMS_200_, TMS_60_, TMS_30_). MEP amplitudes were also labelled according to the muscles from which they were registered (FDI, ADM). EMG onset times were referenced to the ones in the non-TMS trials. A 2-way rmANOVA (TIME x PARADIGM was performed to compare EMG onset times across conditions. A 3-way rmANOVA (TIME x PARADIGM x MUSCLE) was performed to test for changes in MEPs. Post-hoc comparisons were run in the case of finding significant effects. Alpha P-levels obtained from paired comparisons between BASE and the other TMS conditions were Bonferroni-corrected by multiplying them by a factor of 4.

To assess the effect of the TMS on movement times in experiment 2, we used non-TMS trials to estimate the distribution of the lengths of the TMS-to-EMG onset intervals had participants not been biased by the TMS. To do this, all non-TMS trials obtained from each session were used by a simulation algorithm (500 iterations) that, for each participant: *i*) randomly selected 5 trials; *ii*) obtained a simulated TMS time for the “next” trial as in the actual experiment; *iii*) randomly selected a new trial; *iv*) obtained the time interval between the FDI EMG onset time and the simulated TMS time and kept it if it was positive (*i.e.*, TMS delivered before the movement). The resulting simulated distributions of TMS-to-EMG onset intervals were compared with the registered TMS-to-EMG onset intervals by comparing their histograms. This comparison was performed by running individual Wilkoxon tests between bins of 40 ms of width of the two histograms (real and simulated) from 1 s before the EMG onsets. The resulting *P*-values were corrected for multiple comparisons by multiplying them by the number of bins assessed (*i.e.*, 26 bins). This comparison was run separately for TMS-to-EMG onset intervals obtained using the right and left hand FDI muscles to assess their similarity. Additionally, using the TMS trials of all participants, we assessed the difference between the EMG onsets in the left and right hand FDI muscles as a function of the interval between the times at which TMS was delivered in each trial and the corresponding EMG onset time in the left hand FDI muscle. This analysis was done to justify the need of using EMG onsets from the left hand FDI muscle by showing that EMG onsets in the right hand FDI were delayed when TMS pulses were given in close proximity to movement times (Ziemann et al. 1997). This assessment was done using a bootstrapping approach equivalent to the one used to look for changes in MEP amplitudes and explained in the next paragraph.

For the MEP analysis in the SP task in experiment 2, the times of the TMS pulses relative to the subsequent movements could not be well controlled (due to the free nature of SP movements). To overcome this limitation, bootstrap statistics were applied to all participants’ MEP amplitudes in order to identify, in an unbiased way, TMS-to-movement intervals of interest, *i.e.*, intervals between TMS pulses and FDI EMG onsets within which significant increases or decreases in MEP amplitudes were observed. For this analysis, the left FDI EMG onsets were considered since right FDI EMG onsets could be biased by the TMS-induced delays of voluntary actions in the trials where the stimulus was given in close proximity with the subsequent movement (Ziemann et al. 1997). To look for intervals of interest, the following steps were repeated for 100 iterations: 1) 200 TMS-trials per participant were chosen at random; 2) MEPs selected from each participant were referenced to baseline (BASE) MEPs obtained more than 500 ms before the estimated left FDI EMG onset times by computing the z-scores, *i.e.* MEPs were subtracted the mean and divided by the standard deviation of the BASE MEPs; 3) MEP amplitude values from all participants were merged; 4) a sliding window of 40 ms in steps of 20 ms was applied from −1 s to the movement onset. For each window, 40 MEPs were picked at random with replacement and used to calculate a mean (Graimann et al. 2002). This was repeated 1000 times, thus generating 1000 means for every window. The 5^th^ and 995^th^ ranked values were taken as confidence intervals. After this process, an average of all estimated confidence intervals was taken to produce the definitive confidence intervals of MEP changes across the time in preparation for movements. Subsequently, a 2-way rmANOVA with factors TIME and MUSCLE (FDI and ADM) was run to study the changes in MEP amplitudes. The TIME factor included the BASE period (more than 500 ms before movements), and the periods of time in which significant decreases or increases in the right FDI MEPs were observed (referred to as TMSDEC and TMSFAC conditions in the results section). Alpha P-levels obtained from paired comparisons between BASE and the other TMS conditions were Bonferroni-corrected.

Throughout the manuscript, results are reported as group mean ± SEM and *P* values < 0.05 are considered to be significant, unless indicated otherwise. The Greenhouse– Geisser procedure was applied where necessary to correct for violations of sphericity in rmANOVAs.

## Results

### Experiment 1

#### Movement times

The average movement onset times (based on EMG) obtained in the non-TMS trials were −7 ± 6 ms in the PT (indicating that participants successfully synchronised EMG activity to the onset of the trigger) and 211 ± 5 ms in the RT paradigms, respectively. The average EMG onset times in non-TMS trials of the experimental blocks were not significantly different to the ones estimated during the initial training phase prior to the start of the experimental sessions (−19 ± 5 ms, *P* = 0.135 for PT; 206 ± 3 ms, *P* = 0.277 for RT).

Fig. 2 shows the average EMG onset time in TMS trials relative to non-TMS trials; positive values indicate EMG onset is delayed. The TMS pulse appears to produce two effects. First, EMG onset is delayed in both RT and PT trials if the TMS pulse is timed to occur 30 or 60ms prior to EMG onset as determined in non-TMS trials (TMS_30_ and TMS_60_). This effect is caused by two factors: *i*) in TMS trials, movements that are already present before the TMS are not taken into account to estimate movement times, whereas they would have been included in non-TMS trials; and also importantly *ii*) TMS at the intensity used in this study produces an MEP followed by a silent period that delays volitional EMG onset (Ziemann et al. 1997). The second effect of TMS in Fig. 2 is only seen in the RT trials. In this case reaction times are shortened when TMS is given 200 ms prior to average EMG onset times. Since the mean reaction time is 211 ms (see above) this means that reactions are speeded when the TMS pulse is applied at around the time of the imperative stimulus. Previous work refers to this effect on reaction times as a form of IF (Terao et al. 1997). Interestingly, TMS pulses applied at a similar time prior to EMG onset in the PT had no effect on movement time.

Table 1 summarises the main results obtained from the statistical analysis done on movement times. There was a significant main effect of TIME (*F*_[4,11]_ = 12.035; *P* < 0.001) with post-hoc paired comparisons showing significantly delayed EMG onset times for TMS_60_ and TMS_30_ conditions compared to BASE (*P* < 0.001). In addition, there was a significant effect of the PARADIGM x TIME interaction (*F*_[4,11]_ = 8.652; *P* < 0.001) due to a significant difference between RT and PT conditions when the TMS pulse was given 200 ms before mean EMG onset (TMS_200_) (*P* < 0.001). The EMG onset times were significantly reduced in the RT task for TMS_200_ condition compared to BASE (*P* < 0.028). No other significant differences were found between paradigms for EMG onset times (*P* > 0.1 in all cases).

**Table 1.** Results of rmANOVAs on movement times (experiment 1) and MEP amplitudes (experiments 1 and 2)

|  | F[df,error] | P |
| --- | --- | --- |
| **Experiment 1 - Movement Times** |  |  |
| PARADIGM | 3.097[1,14] | 0.100 |
| TIME | 12.035[4,11] | <0.001 |
| PARADIGM x TIME | 8.652[4,11] | <0.001 |

| Experiment 1 - MEP Amplitudes |  |  |
| --- | --- | --- |
| PARADIGM | 0.560<sub>[1,14]</sub> | 0.467 |
| MUSCLE | 20.511<sub>[1,14]</sub> | <0.001 |
| TIME | 6.407<sub>[4,11]</sub> | <0.001 |
| PARADIGM x MUSCLE | 0.016<sub>[1,14]</sub> | 0.900 |
| PARADIGM x TIME | 3.455<sub>[4,11]</sub> | 0.024 |
| MUSCLE x TIME | 7.624<sub>[4,11]</sub> | <0.001 |
| PARADIGM x MUSCLE x TIME | 1.468<sub>[4,11]</sub> | 0.065 |
| Experiment 2 - MEP amplitudes |  |  |
| MUSCLE | 22.154<sub>[1,17]</sub> | <0.001 |
| TIME | 14.940<sub>[2,16]</sub> | 0.001 |
| MUSCLE x TIME | 8.132<sub>[4,11]</sub> | <0.001 |

#### Corticospinal excitability

Resting motor threshold and 1mV levels were 49 ± 3 % and 60 ± 3 % of the maximum stimulator output, respectively. On average, 21 ± 5 MEPs (mean ± SD) were averaged for each TMS condition and subject (averages per TMS condition are shown in Fig. 3). The number of averaged MEPs was comparable across TMS conditions except for TMS_30_ in which a smaller number was used’ since TMS at this timing was more likely to occur after the onset of the EMG, contaminating measurements of the MEP.

**Figure 3.**
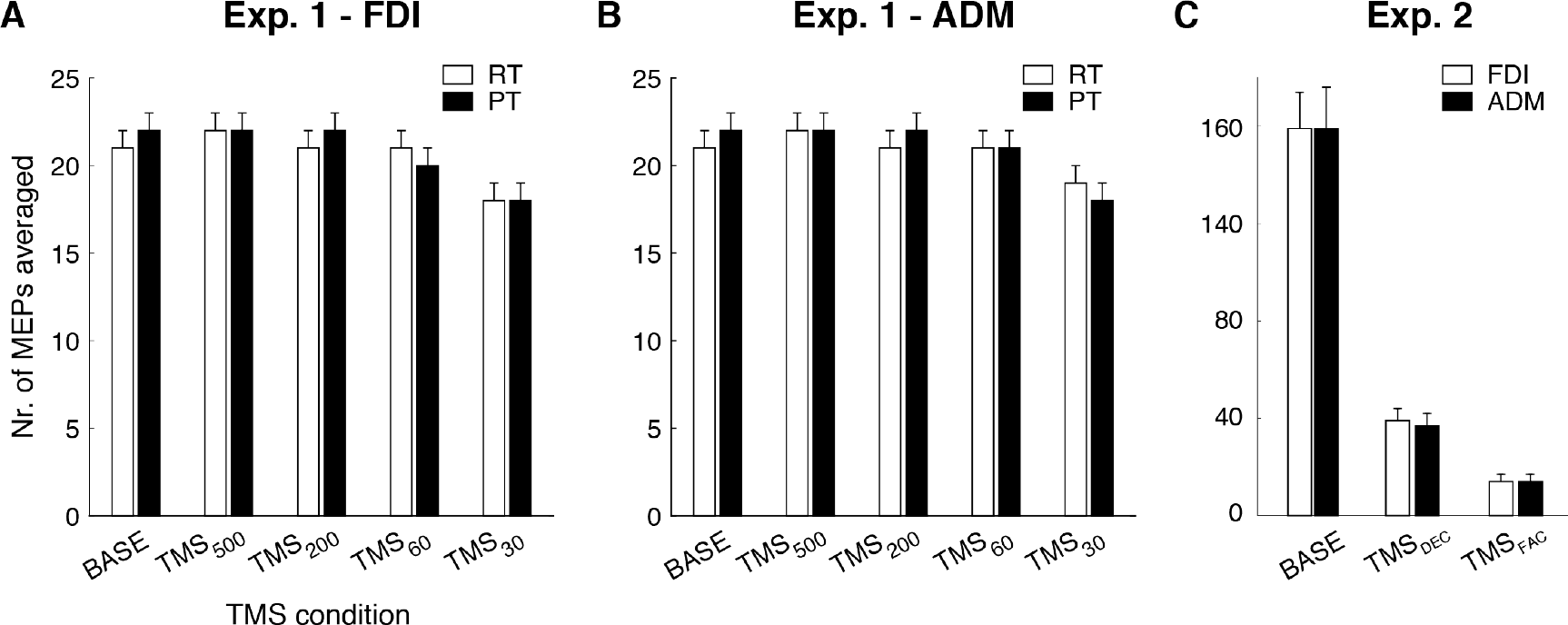
Average number of MEPs considered per muscle, paradigm and TMS condition.

Fig. 4 shows the average amplitude of MEPs evoked at different times relative to mean EMG onset in in both RT and PT tasks. In both tasks, FDI and ADM MEPs were reduced when the TMS pulse was given 200 ms before mean EMG onset (TMS_200_), which is in line with previous studies of preparatory inhibition using RT paradigms (Duque et al. 2010; Lebon et al. 2016; Hannah et al. 2018). TMS pulses delivered 30 and 60 ms prior to EMG onset reflected an increase of CSE in the FDI (agonist muscle) towards the movement initiation time, while MEPs continued to be suppressed in the (task-irrelevant) ADM (Chen and Hallett 1999; Mackinnon and Rothwell 2000).

**Figure 4.**
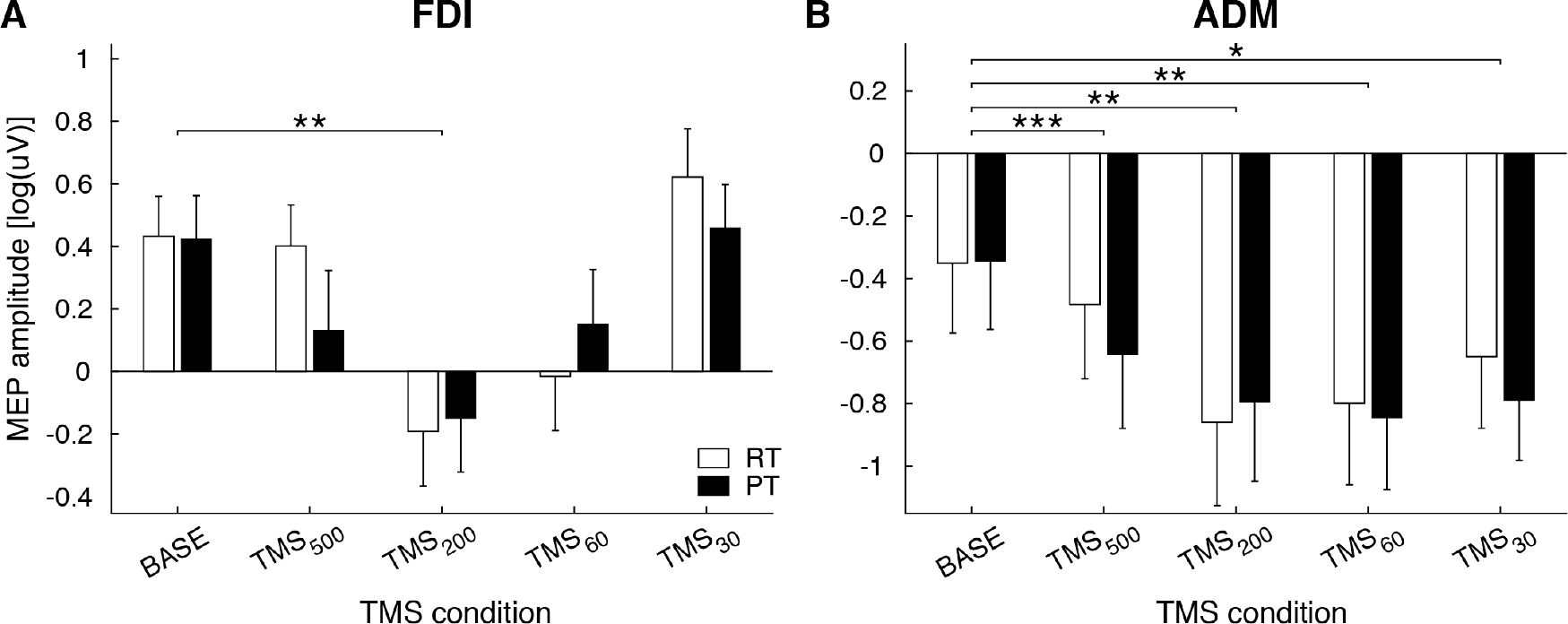
MEP amplitudes in the FDI (A) and ADM (B) before movements made in the RT and PT tasks. Importantly, there was no significant difference between the two paradigms in the task-related muscle at TMS_200_. \**P* < 0.05; ** *P* < 0.01; *** *P* < 0.001, compared with BASE.

Table 1 summarises the statistical results. There were no significant differences between BASE MEP amplitudes in the RT and PT tasks (*P* = 0.467). A three way rmANOVA with main factors of TIME (BASE, TMS_500_, TMS_200_, TMS_60_, TMS_30_), PARADIGM (RT, PT) and MUSCLE (FDI, ADM) revealed significant main effects of TIME (*F*_[4,11]_ = 6.407; *P* = 0.006) and MUSCLE (*F*_[1,14]_ = 20.511; *P* = 0.006). Post-hoc paired comparisons revealed that BASE MEP amplitudes were significantly higher than for TMS_500_ (*P* = 0.016), TMS_200_ (*P* = 0.001) and TMS_60_ (*P* = 0.014) conditions. There was also a significant interaction of MUSCLE x TIME (*F*_[4,11]_ = 7.624; *P* = 0.003). Post-hoc comparisons revealed significant reductions of MEP amplitudes for condition TMS_200_ compared to BASE in the FDI (*P* = 0.003), whereas in the ADM, MEP amplitudes were significantly reduced at many more timings relative to BASE MEPs: TMS_500_ (*P* < 0.001), TMS_200_ (*P* = 0.004), TMS_60_ (*P* = 0.006) and TMS_30_ (*P* = 0.040). Finally, there was a significant interaction of PARADIGM x TIME (*F*_[4,11]_ = 3.455; *P* = 0.033). Post-hoc comparisons revealed significant differences between MEPs in the PT and RT tasks for TMS_500_ (*P* = 0.038), thus suggesting that the suppression of CSE started earlier in the PT task (note that this difference is not shown in Fig. 4, where results are presented separately for each muscle). This was expected since movements in PT trials are performed earlier (relative to the beginning of trials) than movements in RT trials, and therefore, MEP suppression is likely to start earlier as well. Importantly, there was no significant difference in the amount of MEP suppression in the FDI for TMS_200_ condition between the RT and PT paradigms (*P* = 0.630). For TMS_200_, FDI MEPs relative to BASE MEPs showed an average reduction of 41 % (RT task) and 40 % (PT task).

### Experiment 2

#### Movement times

Participants tended to initiate movements about every 5 s (4.94 ± 0.16 s and 4.99 ± 0.18 s in TMS and non-TMS trials respectively). Participants had no difficulties in performing bilaterally synchronous movements: the average (across paricipants) difference in non-TMS trials between the onset of EMG in the right and left FDI was 2 ± 1 ms (individual differences ranged from −17 ms to +14 ms). Fig. 5-A shows how the interval between the onset of right and left FDI EMG varies as a function of the time of the TMS pulse measured relative to the onset of the left EMG. The figure shows that in most trials the hands move synchronously (*i.e.*, a mean difference close to zero). However, when the TMS pulse (to the left hemisphere) occurs closer than 200 ms to EMG onset in the left (“unstimulated”) FDI, the onset of EMG in the right FDI is delayed by up to 50 ms. As in experiment 1, this is because the TMS pulse evokes an MEP in the right hand that is followed by a silent period that delays onset of EMG on that side (Ziemann et al. 1997).

**Figure 5.**
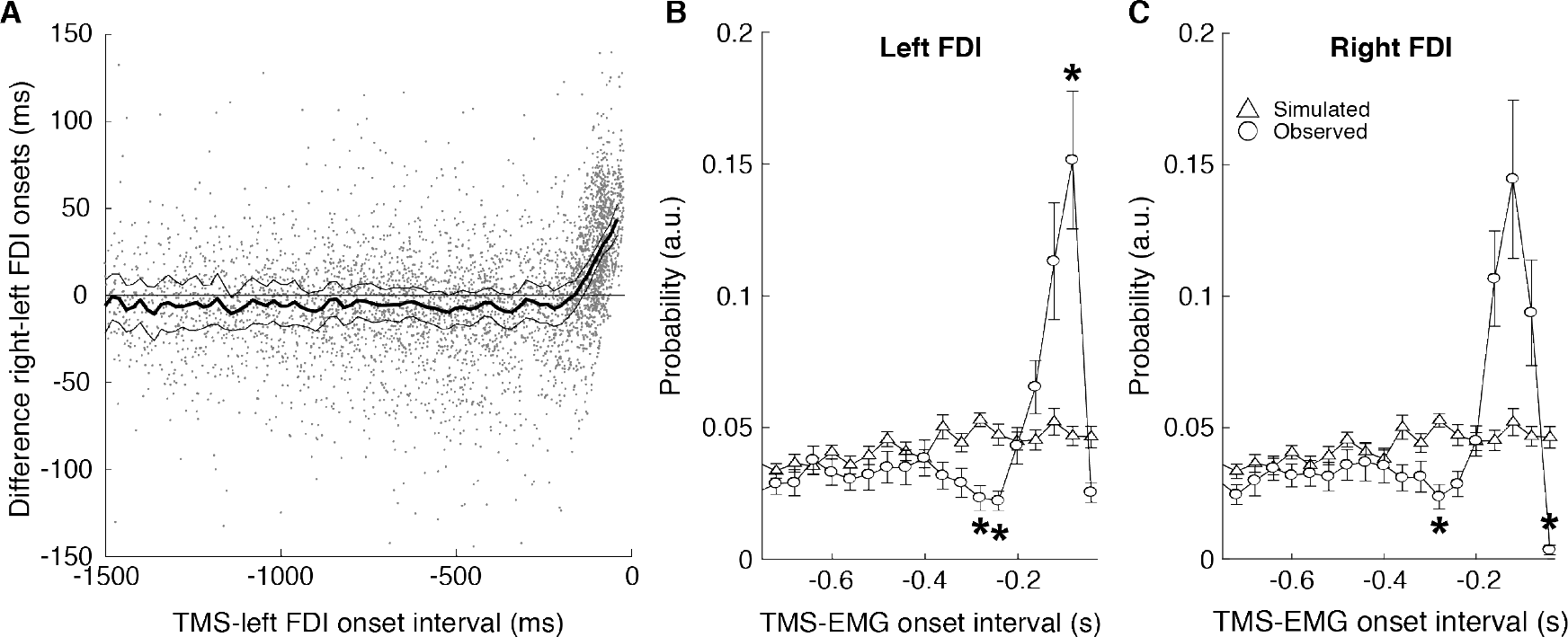
EMG onset times relative to TMS in the SP task. (A) Intervals between left and right FDI EMG onsets as a function of the interval existing between the TMS pulses and the movement time estimated using the left hand FDI EMG onset in the SP task in experiment 2. Dots represent individual trials of all participants. The black traces indicate the average and upper and lower limits (*P* < 0.01) of the estimates of the mean differences between the left and right FDI EMG onsets computed in sliding windows of 40 ms. The traces show an effect of the TMS on right FDI onsets when stimuli are delivered less than 200 ms before the estimated EMG onset. (B-C) Real (circles) and simulated (triangles) intervals between the TMS pulses and posterior movements in TMS trials. Traces represent the observed distributions of TMS-to-EMG onset intervals for the left (B) and right (C) FDI muscles. Graphs are obtained by combining data from all subjects. \**P* < 0.05, TMS-to-EMG onset intervals where a significant difference across participants exists between the simulated and observed data.

Figs. 5-B and 5-C show the probability of a TMS pulse being delivered at different times prior to EMG onset on the left and right sides. The two sets of symbols plot the observed distribution and the distribution calculated by assuming that the time of the TMS pulse is independent of the onset of EMG. Compared to the simulated distribution, the observed distribution of TMS-to-EMG onset probabilities has a trough around 280-260 ms (Wilcoxon test: *P* = 0. 011/0.015−280/260 ms- and *P* = 0.018−280 ms- for the left and right hand FDIs, respectively) followed by a peak at 80 ms before the EMG onset (*P* = 0. 029 for the left hand FDI). In other words, there are fewer trials than expected in which a movement starts around 280-260 ms after a TMS pulse. Conversely there are more trials than expected in which EMG onset occurs around 80 ms after a TMS pulse. This suggests that, in trials in which TMS was delivered around 280-260 ms before participants were about to move, button presses were performed earlier than they would have been, with the result that EMG onsets are not independent of the time of the TMS pulse. Effectively, a TMS pulse given 280-260 ms before an intended movement advances movement onset so that we rarely observe TMS at this interval. Unfortunately, it is difficult to estimate by how much the onset is advanced. For example, if the EMG onset is advanced by 80 ms (Smith et al. 2019), we might have expected to see an increase in the number of trials at 200 ms interval. However, TMS pulses at 200 ms might also advance movement onset, in which case the additional counts from speeded trials at −280 ms would not be observed etc. Note that this effect is seen in both left and right FDI muscles, and we suggest in the discussion that it is related to the phenomenon known as intersensory facilitation (IF), which is typically studied using RT movements.

#### Corticospinal excitability

Resting motor threshold and 1 mV levels were 56 ± 3 % and 66 ± 4 % of the maximum stimulator output. Fig. 3 (right) summarizes the average number of FDI and ADM MEPs used to characterize excitability states in the three TMS conditions considered.

Fig. 6 shows the summary of the MEP results obtained both using bootstrap statistics on the grouped (z-scored) data and from the posterior comparison between the intervals of interest. The bootstrap analysis in fig. 6A-B indicates that in the FDI (agonist muscle) there is a period around −180:−100 ms during which the MEP was significantly smaller than at baseline (both confident intervals below 0). This disappears at around −80 ms, after which it is followed by a final period of facilitation. The reduction is also seen in (non-involved) ADM over a similar time period. We defined a period of reduced excitability (TMS_DEC_) from −180 ms to −100 ms. A late premovement phase was defined as the final 80 ms interval before EMG onset to have large enough set of samples of valid MEP amplitudes to average for that condition (TMS_FAC_) (Pascual-Leone, Valls-Sole, et al. 1992; Chen and Hallett 1999; Chen et al. 1999). The main results of the rmANOVA test comparing BASE, TMS_DEC_ and TMS_FAC_ are summarized in Table 1. In line with the results obtained in experiment 1, there was a significant main effect of TIME (*F*_[2,16]_ = 12.544; *P* = 0.001). The post-hoc comparison between MEPs for BASE and TMS_DEC_ conditions revealed a significant reduction of MEP amplitudes at the latter time point (*P* = 0.001). There was also a significant interaction of MUSCLE x TIME (*F*_[2,16]_ = 12.733; *P* = 0.002). Post-hoc comparisons revealed significant differences between the FDI MEPs recorded in the intervals BASE and TMS_DEC_ (*P* = 0.002). In the ADM, comparisons revealed significant reductions (relative to BASE) of MEPs at TMS_DEC_ (*P* = 0.006) and TMS_FAC_ (*P* = 0.018).

**Figure 6.**
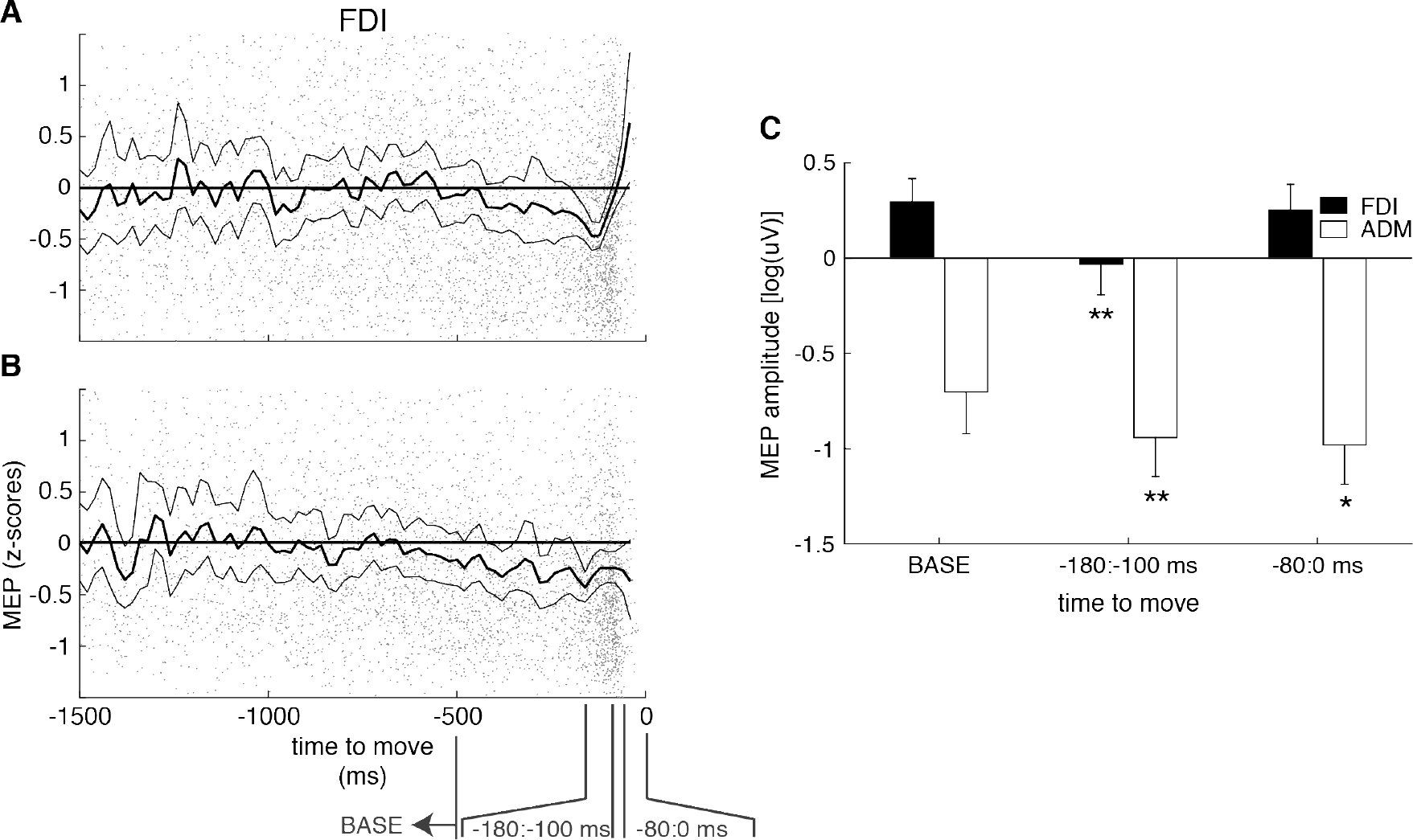
Changes in the FDI (A) and ADM (B) MEPs across time before movements in the SP task. Dots (in grey) show z-scores of MEP amplitudes (all participants and trials). Solid traces represent the upper and lower confidence limits (thin traces) and the mean of the observations at each point in time (thick trace), both obtained using a bootstrap analysis with all data points (*P* < 0.01). Marks at the bottom identify three intervals of interest used to extract MEPs for a subsequent group level rmANOVA test. These are aimed to contain BASE MEPs, and MEPs during the periods of reduced excitability (TMS_DEC_) and movement initiation phase (TMS_FAC_). (C) Normalized group means and SEs of FDI and ADM MEP amplitudes in the three intervals of interest and TMS conditions where significant changes in MEPs are found. \**P* < 0.05; ** *P* < 0.01.

## Discussion

In the present experiments, we probed the temporal evolution of CSE in paradigms with different constraints on movement initiation times. The results showed that reduced excitability can be observed prior to a voluntary movement whether it is self-paced, predictive or reactive. It therefore seems unlikely, particularly in the self-paced task, that preparatory corticospinal inhibition is necessary to prevent premature release of movement (“impulse control”). The results are more consistent with hypotheses that view corticospinal inhibition as an essential part of movement preparation. Unexpectedly, our data also revealed that the TMS pulse had distinct biasing effects on movement onset times across the different paradigms. While the initiation of a predictive movement was unaffected by TMS, RT and SP movements were speeded in a way resembling IF effects commonly reported in RT tasks (Nickerson 1973). We propose that this reveals similarities in the way planned voluntary movements are triggered in the presence or absence of specific external cues.

### A period of reduced corticospinal excitability precedes EMG onset in all three types of movement

CS excitability was reduced for a short period about 200 ms prior to EMG onset in RT and PT tasks, and about 140 ms prior to EMG onset in the self-paced task. In the RT task this corresponded to the approximate time of the imperative signal. Overall, the size and spatio-temporal patterns of changes in CSE were very similar to earlier work. On average, MEP amplitudes decreased by 35%, in line with previous data (Duque et al. 2014; Quoilin et al. 2016), and MEP suppression was only maintained in the task-unrelated muscle (the ADM in our case).

The three hypotheses (see Introduction) put forwards to account for preparatory CS suppression were developed during the study of RT movements. One of these (“competition resolution”) is not relevant to the present study since the same movement was performed on each trial, such that there was no need to suppress possible competing responses. As such we confine the discussion to the “impulse control” and “spotlight” hypotheses.

The “impulse control” theory proposes that in RT tasks, reduced CSE is a mechanism that reduces the probability of premature release of the prepared movement (Duque and Ivry 2009; Duque et al. 2010, 2012). However, this is not a satisfactory explanation for the very similar effect in the self-paced task since there is explicitly no temporal constraint on initiation time, and no chance of premature release. Strictly speaking, suppression is also unnecessary in the PT movements since, if preparation was started at the correct time, it would automatically evolve to reach threshold and initiate a movement coinciding with the external event. Finally, the idea that MEPs during movement preparation reflect a process of proactive inhibition is also at odds with the observation that, although we observed similar suppression of CSE in both RT and PT tasks, there were marked differences in the way the TMS pulses interfered with movement onset times (i.e. intersensory facilitation (IF) was only seen in RT). Although it is possible that reduced CSE has different functions in different types of movement, it seems more likely that it reflects a process common to preparation of all movement types.

An alternative interpretation for suppression of CSE that could apply equally to all three movement types is the “spotlight” hypothesis that movement selection is facilitated during corticospinal suppression because detection of incoming excitatory input is easier against a quiescent background (Hasbroucq et al. 1997; Greenhouse et al. 2015; Duque et al. 2017). Although this could account for our present results, as noted in the Introduction, it does not readily explain the observed selectivity in the suppression levels observed in different M1 circuits projecting to the CS tract (Hannah et al. 2018). Instead, our results appear a better fit for the alternative hypothesis that changes in CSE reflect properties of neural populations evolving towards more stable states from which to initiate movement (Churchland et al. 2010; Shenoy et al. 2013; Kaufman et al. 2014, 2016; Hannah et al. 2018). In fact, recent studies in primates show that movements initiated by different types of triggers are all preceded by the same patterns of activity during which discharge rates change without any overt EMG activity (Lara et al. 2018). The time spent in this preparatory state can be compressed or extended depending on task demands (Lebon et al. 2016; Lara et al. 2018). This observation might explain why MEP suppression can be modulated by the duration of the preparatory period (Lebon et al. 2016). It may also be relevant to choice reaction tasks without prior warning cues in which MEPs remain unchanged before the imperative signal but show a suppression right after the onset of the imperative cue (Duque et al. 2014). Finally, it may account for the later timing of suppression in SP movements: the lack of constraint on precise movement onset in self-paced movements may allow a faster transition through preparatory states.

We cannot comment on the location of this suppression since the TMS pulse recruits activity in neural populations of cortex, brainstem and spinal cord, and it is likely that preparation proceeds in parallel at all levels of the motor output. Nevertheless, the conclusion is that preparatory suppression of CSE is an integral part of movement initiation, irrespective of the way the movement is triggered.

### Intersensory facilitation is absent in predictive tasks

In addition to evoking MEPs, the TMS pulse could also affect the timing of the volitional motor response: in both RT and PT movements, pulses timed within about 50 ms of the expected EMG onset delayed movement. Previous workers have ascribed this effect to the silent period following the MEP, which suppresses EMG onset (Ziemann et al. 1997). In addition to this delaying effect, in RT only, pulses that occurred earlier, around the time of the imperative stimulus to move, speeded up EMG onset. The effect is thought to be due to the sensory input produced by the TMS pulse (the “click” of the coil plus stimulation of the skin and muscle of the scalp) and has been interpreted as intersensory facilitation (IF) (Nickerson 1973; Terao et al. 1997). The temporal conjunction of the imperative stimulus with the additional sensory input from the TMS pulse is thought to shorten the time for identification of the go-signal and speed up the onset of EMG (Pascual-Leone, Valls-Sole, et al. 1992).

This speeding effect was not present in the predictive task, even when the TMS pulse occurred at a similar time with respect to the onset of EMG. The implication is that predictive movements are triggered differently to RT movements. In eye movement studies, saccades triggered by predictably timed imperative stimuli are thought to employ internal mechanisms that anticipate the timing of the external stimulus (Janssen and Shadlen 2005; Badler and Heinen 2006). It may be that similar internal timing mechanisms are used in our predictive task, and these trigger the release of movement so that it coincides with timing of the last countdown cue. In this case, external signals during the delay period of PT movements may be either downregulated (Alink et al. 2010) or simply disregarded (Rohenkohl et al. 2012), with the result that IF is absent.

### TMS biases movement onset in self-paced movements

Analysing brain responses to external stimuli before SP movements with a degree of temporal precision is technically challenging and has only been attempted in a few studies. The work most relevant to the present results is that of Castellote and colleagues who showed that StartReact responses (*i.e.*, speeded responses caused by a startling stimulus) prior to self-initiated movements are comparable to StartReact responses in RT paradigms (Valls-Sole et al. 1999; Castellote et al. 2013). Interestingly, Castellote’s experiment showed a biasing effect of startling stimuli on movement times that closely resembles that obtained here (Fig. 5), *i.e.*, if delivered approximately 300 ms before the forthcoming movement, startling stimuli speed-up movement onset. The features of the responses matched those obtained in StartReact paradigms using RT tasks, which allowed the authors to suggest that mechanisms engaged in the preparation for SP and cue-driven actions shared common elements (Castellote et al. 2013). In our case, the intensity of the applied stimuli (1 mV TMS and participants using ear defenders, which lessen the likelihood of startle response) suggests that the observed effects are closer to IF (Pascual-Leone, Brasil-neto, et al. 1992; Pascual-Leone, Valls-Sole, et al. 1992). Quantifying precisely how much movements are sped up would help verify this idea (Valls-Sole et al. 2008), but doing so is challenging because of the lack of a more precise knowledge about when movements would be performed in the absence of external stimuli. Based on the fact that the stimuli used in RT and SP paradigms were equal and responses to TMS comparable, it is conceivable that effects observed in both cases reflect the use of a similar neural strategy to trigger actions that is not shared by PT movements.

### Technical considerations and future work

Studying CS excitability changes before SP movements with TMS has inherent limitations. We were able to obtain movement onset times and MEPs in our SP paradigm despite the apparent difficulties in accurately probing excitability at specific, well-defined times relative to movement onset. To achieve this, participants were asked to perform bimanual movements, so that muscle activity in the non-stimulated hand could be used to estimate the EMG onset times. This differed from the unilateral movements used in experiment 1. Importantly, this difference is not expected to have any influence on the obtained results according to previous research showing comparable MEP suppression in unimanual and bimanual movements (Duque and Ivry 2009). Additionally, in experiment 2, the estimations of TMS times (relative to the movements) were done based on the times of the subsequent movements of the non-stimulated hand, which also presents IF effects, is not affected by cortical silent period-related delaying effects (Ziemann et al. 1997), but may still have been biased by the TMS in a different way than the stimulated side. Therefore, precise estimations of the excitability reduction peak time in this case are not definitive.

Previous studies testing the possible contribution of spinal mechanisms to the decrease seen in MEP amplitudes in preparation for movements in RT tasks have led to contradictory results (Duque et al. 2010; Lebon et al. 2016; Hannah et al. 2018). The techniques used here to probe CSE do not allow us to separate out the contribution of spinal inhibitory processes to the observed MEP changes. Knowing if and how spinal inhibition applies to cue-driven and SP movements could help further to test the hypothesis that observed preparatory changes are not driven by proactive control mechanisms.

Finally, we did not use catch trials to make the RT and PT tasks as similar as possible (we could not use catch trials in the PT task because otherwise movements would have been initiated in a reactive way after knowing whether a movement was required in each trial). Participants were instructed to use two well differentiated strategies to trigger their movements. In the PT task, participants had to learn how to synchronize their movement with that of the circles. In the RT task, participants were specifically instructed to react to the “GO” instruction given by the overlap of the four circles at the intersection point of the cross. None of our participants reported having difficulties in avoiding the prediction of the “GO” instruction in the RT task. The posterior analysis of the obtained results corroborates this statement: reaction times are in the range of what the literature reports and, when TMS was applied, RT movements (and not PT movements) showed a clear IF effect, which is a distinctive feature of preparatory states in RT tasks extensively described in the literature (Nickerson 1973; Terao et al. 1997; Hannah et al. 2018).

### Conclusions

CSE, as assessed using TMS, is transiently suppressed prior to initiation of RT, PT and SP movements. This may indicate that it represents a common state through which neural activity must evolve in the transition from rest to movement. It may be related to mechanisms that maintain constant corticospinal output at a time when preparatory neural activity in the cortex undergoes rapid change. In addition to probing CSE, the TMS pulse also produces a strong sensory input. In RT movements, this results in intersensory facilitation that speeds up onset of movement. A similar effect is seen in SP movements, suggesting the existence of common neural mechanisms to trigger movement onset. This effect is not present in PT tasks, implying they are triggered differently, perhaps because priority is given to internal signals that predict the time of movement onset.

## ACKNOWLEDGEMENTS

We gratefully acknowledge the technical assistance of Paul Hammond. We also thank Arisa Reka for assistance in collection of data in Experiment 2.

## References

Alink A, Schwiedrzik CM, Kohler A, Singer W, Muckli L. 2010. Stimulus Predictability Reduces Responses in Primary Visual Cortex. J Neurosci. 30:2960–2966.

Badler JB, Heinen SJ. 2006. Anticipatory Movement Timing Using Prediction and External Cues. J Neurosci. 26:4519–4525.

Bestmann S, Duque J. 2015. Transcranial Magnetic Stimulation: Decomposing the Processes Underlying Action Preparation. Neurosci. 1–14.

Burle B, Vidal F, Tandonnet C, Hasbroucq T. 2004. Physiological evidence for response inhibition in choice reaction time tasks. Brain Cogn. 153–164.

Castellote JM, Van Den Berg MEL, Valls-Sole J. 2013. The startreact effect on self-initiated movements. Biomed Res Int. 2013.

Chen R, Corwell B, Hallett M. 1999. Modulation of motor cortex excitability by median nerve and digit stimulation. Exp Brain Res. 129:77–86.

Chen R, Hallett M. 1999. The time course of changes in motor cortex excitability associated with voluntary movement. Can J Neurol Sci. 26:163–169.

Churchland MM, Cunningham JP, Kaufman MT, Ryu SI, Shenoy K V. 2010. Cortical Preparatory Activity: Representation of Movement or First Cog in a Dynamical Machine? Neuron. 68:387–400.

Duque J, Greenhouse I, Labruna L, Ivry RB. 2017. Physiological Markers of Motor Inhibition during Human Behavior. Trends Neurosci. 40:219–236.

Duque J, Ivry RB. 2009. Role of corticospinal suppression during motor preparation. Cereb Cortex. 19:2013–2024.

Duque J, Labruna L, Cazares C, Ivry RB. 2014. Dissociating the influence of response selection and task anticipation on corticospinal suppression during response preparation. Neuropsychologia. 65:287–296.

Duque J, Labruna L, Verset S, Olivier E, Ivry RB. 2012. Dissociating the Role of Prefrontal and Premotor Cortices in Controlling Inhibitory Mechanisms during Motor Preparation. J Neurosci. 32:806–816.

Duque J, Lew D, Mazzocchio R, Olivier E, Ivry RB, Louvain D, Brussels B-, Clinica N. 2010. Evidence for Two Concurrent Inhibitory Mechanisms during Response Preparation. J Neurosci. 30:3793–3802.

Graimann B, Huggins JE, Levine SP, Pfurtscheller G. 2002. Visualization of significant ERD/ERS patterns in multichannel EEG and ECoG data. Clin Neurophysiol. 113:43–47.

Greenhouse I, Sias A, Labruna L, Ivry RB. 2015. Nonspecific Inhibition of the Motor System during Response Preparation. J Neurosci. 35:10675–10684.

Hamada M, Galea JM, Di Lazzaro V, Mazzone P, Ziemann U, Rothwell JC. 2014. Two Distinct Interneuron Circuits in Human Motor Cortex Are Linked to Different Subsets of Physiological and Behavioral Plasticity. J Neurosci. 34:12837–12849.

Hannah R, Cavanagh XSE, Tremblay S, Simeoni S, Rothwell JC. 2018. Selective suppression of local interneuron circuits in human motor cortex contributes to movement preparation. J Neurosci. 38:2869–17.

Hannah R, Rothwell JC. 2017. Pulse Duration as Well as Current Direction Determines the Specificity of Transcranial Magnetic Stimulation of Motor Cortex during Contraction. Brain Stimul. 10:106–115.

Hasbroucq T, Kaneko H, Akamatsu M, Possamaı̈ C-A. 1997. Preparatory inhibition of cortico-spinal excitability: a transcranial magnetic stimulation study in man. Cogn Brain Res. 5:185–192.

Janssen P, Shadlen MN. 2005. A representation of the hazard rate of elapsed time in macaque area LIP. Nat Neurosci. 8:234–241.

Kaufman MT, Churchland MM, Ryu SI, Shenoy K V. 2014. Cortical activity in the null space: Permitting preparation without movement. Nat Neurosci. 17:440–448.

Kaufman MT, Seely JS, Sussillo D, Ryu SI, Shenoy K V., Churchland MM. 2016. The Largest Response Component in the Motor Cortex Reflects Movement Timing but Not Movement Type. eNeuro. 3.

Kim H-Y. 2013. Statistical notes for clinical researchers: assessing normal distribution (2) using skewness and kurtosis. Restor Dent Endod. 38:52.

Lara AH, Elsayed GF, Zimnik AJ, Cunningham JP, Churchland MM. 2018. Conservation of preparatory neural events in monkey motor cortex regardless of how movement is initiated. Elife. 7:7:e31826.

Lebon F, Greenhouse I, Labruna L, Vanderschelden B, Papaxanthis C, Ivry RB. 2016. Influence of Delay Period Duration on Inhibitory Processes for Response Preparation. Cereb Cortex. 26:2461–2470.

Lebon F, Ruffino C, Greenhouse I, Labruna L, Ivry RB, Papaxanthis C. 2018. The Neural Specificity of Movement Preparation During Actual and Imagined Movements. Cereb Cortex. 1–12.

Mackinnon CD, Rothwell JC. 2000. Time-varying changes in corticospinal excitability accompanying the triphasic EMG pattern in humans. 633–645.

Nickerson RS. 1973. Intersensory facilitation of reaction time: Energy summation or preparation enhancement? Psychol Rev. 80:489–509.

Pascual-Leone A, Brasil-neto J, Valls-Sole J, Cohen LG, Hallett M. 1992. Simple reaction time to focal transcranial magnetic stimulation. Brain. 115:109–122.

Pascual-leone A, Valls-Sole J, Wassermann EM, Brasil-Neto J, Cohen LG, Hallett M. 1992. Effects of Focal Transcranial Magnetic Stimula Tion on Simple Reaction Time To Acoustic, Visual and Somatosensory. Brain. 115:1045–1059.

Pascual-Leone A, Valls-Sole J, Wassermann EM, Brasil-neto J, Cohen LG, Hallett M. 1992. Effects of Focal Transcranial Magnetic Stimulation on Simple Reaction Time To Acoustic, Visual and Somatosensory stimuli. Brain. 115:1045–1059.

Quoilin C, Fievez F, Duque J. 2019. Neuropsychologia Preparatory inhibition: Impact of choice in reaction time tasks. Neuropsychologia. 129:212–222.

Quoilin C, Lambert J, Jacob B, Klein PA, Duque J. 2016. Comparison of motor inhibition in variants of the instructed-delay choice reaction time task. PLoS One. 11:1–16.

Rohenkohl G, Cravo AM, Wyart V, Nobre AC. 2012. Temporal Expectation Improves the Quality of Sensory Information. J Neurosci. 32:8424–8428.

Rossi S, Hallett M, Rossini P, Pascual-Leone A. 2011. Screening questionnaire before TMS: An update. Clin Neurophysiol. 122:1686.

Schneider C. 2004. Timing of cortical excitability changes during the reaction time of movements superimposed on tonic motor activity. J Appl Physiol. 97:2220–2227.

Shenoy K V., Sahani M, Churchland MM. 2013. Cortical Control of Arm Movements: A Dynamical Systems Perspective. Annu Rev Neurosci. 36:337–359.

Smith V, Maslovat D, Drummond M, Carlsen AN. 2019. A Timeline of Motor Preparatory State Prior to Response Initiation: Evidence from Startle. Neuroscience. 397:80–93.

Terao Y, Ugawa Y, Suzuki M, Sakai K, Hanajima R, Gemba-Shimizu K, Kanazawa I. 1997. Shortening of simple reaction time by peripheral electrical and submotor-threshold magnetic cortical stimulation. Exp Brain Res. 115:541–545.

Touge T, Taylor JL, Rothwell JC. 1998. Reduced excitability of the cortico-spinal system during the warning period of a reaction time task. Electroencephalogr Clin Neurophysiol. 109:489–495.

Valls-Sole J, Kumru H, Kofler M. 2008. Interaction between startle and voluntary reactions in humans. Exp Brain Res. 187:497–507.

Valls-Sole J, Rothwell JC, Goulart F, Cossu G, Muñoz E. 1999. Patterned ballistic movements triggered by a startle in healthy humans. J Physiol. 516:931–938.

Ziemann U, Tergau F, Netz J, Hömberg V. 1997. Delay in simple reaction time after focal transcranial magnetic stimulation of the human brain occurs at the final motor output stage. Brain Res. 744:32–40.

